# Quantifying stem cell derived islet graft volume and composition with [^18^F]F-DBCO-exendin and [^18^F]FDOPA positron emission tomography

**DOI:** 10.1101/2023.12.18.572141

**Authors:** Väinö Lithovius, Salla Lahdenpohja, Hazem Ibrahim, Jonna Saarimäki-Vire, Lotta Uusitalo, Hossam Montaser, Kirsi Mikkola, Cheng-Bin Yim, Thomas Keller, Johan Rajander, Diego Balboa, Tom Barsby, Olof Solin, Pirjo Nuutila, Tove J. Grönroos, Timo Otonkoski

## Abstract

Stem cell derived islets (SC-islets) are being developed as a novel source of beta cells that would enable large scale cell replacement therapy for insulin dependent diabetes. Therapeutic use of SC-islets carries an inherent risk of unwanted growth; and multiple strategies are being explored for optimizing long-term SC-islet graft effectiveness. However, a method for noninvasive *in vivo* monitoring for SC-islet graft safety and efficacy is lacking, as current insulin secretion measurements are inadequate. Here, we demonstrate the potential of positron emission tomography (PET) for monitoring SC-islet grafts using two tracers: GLP1-receptor binding [^18^F]F-DBCO-exendin and dopamine precursor [^18^F]FDOPA. We could detect and longitudinally monitor human SC-islet grafts in calf muscles of immunocompromised mice. Importantly, graft volume quantified with PET strongly correlated with actual graft volume (*r*^2^=0.91 for [^18^F]F-DBCO-exendin). PET using [^18^F]F-DBCO-exendin allowed delineation of cystic structures and its uptake correlated with graft beta cell proportion, enabling study of SC-islet graft purity noninvasively. [^18^F]FDOPA performed similarly to [^18^F]F-DBCO-exendin, but with slightly weaker sensitivity. Uptake of neither tracer was biased in SC-islet grafts genetically rendered hyper- or hypoactive. Insulin secretion measurements under fasted, glucose-stimulated or hypoglycemic conditions did not correlate with graft volume. In conclusion, [^18^F]F-DBCO-exendin and [^18^F]FDOPA PET constitute powerful approaches to noninvasively assess SC-islet graft volume and composition regardless of their functionality. PET imaging could therefore be leveraged for optimizing safety and effectiveness of SC-islet grafts in patients with insulin dependent diabetes.

## INTRODUCTION

Islet or pancreas transplantation remains the only route to independence from exogenous insulin for patients with type 1 diabetes (T1D). One of the major limitations of islet transplantation is the scarcity of cadaveric islets since at least two donors are required for each patient with many requiring a re-transplantation (*1*). Functional islets differentiated *in vitro* from pluripotent stem cells (SC-islets) are being developed to supplement cadaveric islets (*2–5*). In contrast to cadaveric islets, SC-islets are abundant, functionally consistent and can feasibly be genome edited or encapsulated circumventing the need for systemic immunosuppression (*6, 7*). Given these advantages, SC-islet based cell replacement therapy has the potential to revolutionize the treatment of T1D (*8*). SC-islet based cell replacement therapy has already resulted in insulin independence in a few T1D patients after intraportal SC-islet implantation (clinical trial *NCT04786262)* (*9*), with additional trials ongoing (*NCT03163511*, *NCT05791201*).

Nonetheless, clinical use of stem cell derived tissue has inherent risks. Nondirected *in vivo* differentiation of lingering progenitor cells can lead to expansion of undesired pancreatic ductal cysts (*10–16*), potentially hampering therapeutic effectiveness. Furthermore, *in vitro* culture conditions have been shown to enrich chromosomal abnormalities and oncogenic mutations in pluripotent stem cells (*17, 18*). Upon neoplastic expansion, the SC-islet graft can be eliminated by cessation of immunosuppressants and subsequent immune rejection, unless the graft is autologous or immune-evasive. Immune evasion by HLA knockout or overexpression of PD-L1 or CD47 has been utilized for SC-islet implantation studies (*4, 19–23*), but would enable any neoplasm originating from the edited SC-islet graft to readily invade and metastasize (*6, 23, 24*). An extreme demonstration of these risks was recently reported when implantation of autologous stem cell derived tissue claimed to have been differentiated into SC-islets developed a treatment resistant, metastatic teratoma in two months (*25*).

Given both the enormous promise and the potentially substantial risks presented by SC-islet cell replacement therapy, a robust noninvasive method for monitoring the SC-islet grafts *in vivo* is required. The optimal monitoring method would: a) quantify total graft volume b) inform about graft composition such as its beta cell mass and c) not be biased by the beta cells’ functional state. Monitoring graft growth and composition could help assess the *in vivo* impact of updated differentiation protocols, alternative implantation sites and stress-reduction strategies (*22, 26–31*). In terms of safety, detecting excessive increase in graft size or change in its composition could inform immunosuppression cessation or activation of suicide switches (*32, 33*).

Currently, to monitor graft composition it must be removed, preventing longitudinal monitoring. Noninvasive monitoring methods are based on blood C-peptide concentration or magnetic resonance imaging (MRI). C-peptide level is mostly related to beta cell functionality, giving no information about the overall volume of grafts (*34, 35*). When combined with a contrast agent, MRI allows detection of intrahepatic islet grafts in human (*36, 37*) but gives no information about graft composition.

Positron emission tomography (PET) is a functional imaging technique that employs target specific radiotracers for visualizing biochemical and physiological processes *in vivo*, as the most sensitive modality for noninvasive imaging in human. PET and other nuclear imaging methods have been explored in the context of islet transplantation, detecting islet grafts in human muscle (*38*) and liver (*39*) and quantifying graft volume in mouse (*40, 41*). These studies used radiolabeled exendin, targeting the GLP1-receptor, expressed in human primarily in pancreatic beta cells (*42, 43*). In light of this, we recently established ^18^F-labeling of exendin-4 ([^18^F]F-DBCO-exendin-4, referred to as “[^18^F]exendin” throughout this article) (*44*). Imaging the pancreas with ^18^F-labeled DOPA ([^18^F]FDOPA), a dopamine precursor analog, has an established clinical application in the diagnosis of the focal form of congenital hyperinsulinism (CHI)(*45*). Focal CHI presents with a single adenomatous lesion with hypersecretory beta cells, which has a high uptake of [^18^F]FDOPA, while beta cells in normal islets are dormant. This separation allows precise surgical treatment (*46*). Furthermore, a proof-of-concept transplantation study demonstrated that [^18^F]DOPA uptake detects human islet xenografts in mice (*47*).

In this study, we establish the use of PET as a noninvasive monitoring method for human SC-islet grafts. Specifically, we investigate the relationship of the SC-islet graft volume, composition and functional state to the uptake of [^18^F]exendin and [^18^F]FDOPA. We show that [^18^F]exendin and [^18^F]FDOPA can accurately quantify the actual volume of the SC-islet grafts, as well as inform about graft composition.

## RESULTS

### Study setup and validation of tracer receptor expression in SC-islet grafts

We implanted human SC-islets into calf muscles of immunocompromised mice in two cohorts (Fig. 1A). We implanted the first (cohort-1) with different volumes of SC-islets to dissect the relationship between PET tracer uptake and graft volume; and the second (cohort-2) with control SC-islets or genetically engineered hyper- or hypoactive SC-islets, to study the relationship between tracer uptake and graft functional state. Dictated by the differentiation protocols available at the time, the SC-islets used for cohort-1 were immature (*48*), while cohort-2 SC-islets were differentiated with a protocol yielding SC-islets with primary islet - like function and cytoarchitecture (*2, 49*). We followed both cohorts for 5 months with PET/CT imaging using two tracers: [^18^F]exendin, hypothesized to be suited for graft volume quantification; and [^18^F]FDOPA, hypothesized to allow discerning graft functionality (Fig. 1B).

**Figure 1.**
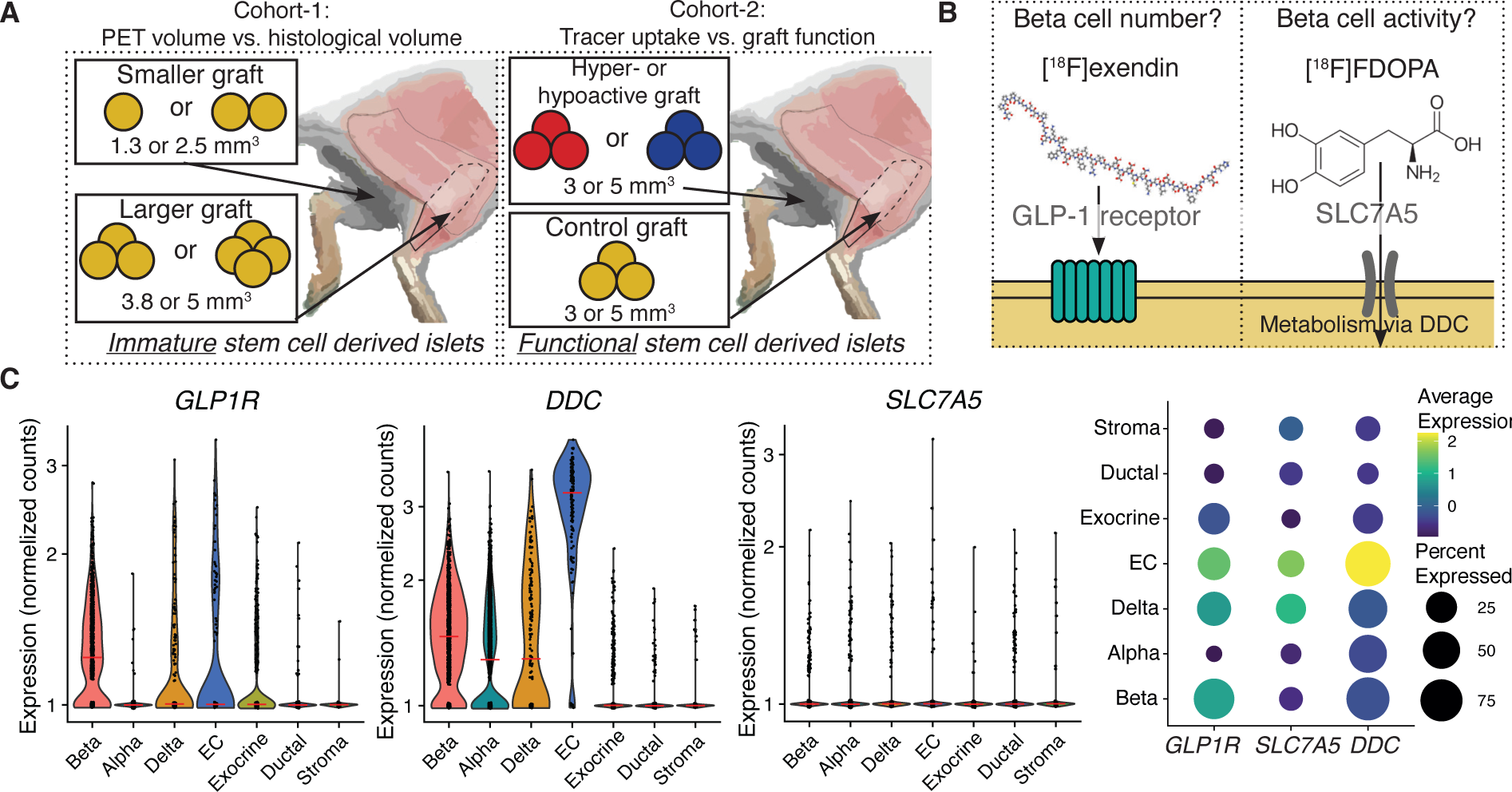
study setup and tracer cell type specificity: **A)** Study setup, cohort-1: comparison of different-sized SC-islet grafts in each leg, cohort-2 consisted of comparison of SC-islet grafts of different functional phenotypes in each leg. Immature SC-islets differentiated as in *Balboa et al. 2018 Elife*, functional ones as in *Barsby et al. 2022 STAR protocols*. 1 mm^3^ = 566 IEQ ≈ 150 SC-islets. **B**) PET-tracers used in the study, [^18^F]F-DBCO-Exendin-4 ([^18^F]exendin) and [^18^F]FDOPA **C**) Tracer target expression in different cell types of control SC-islet kidney subcapsular grafts at 6 months post-implantation, from *Balboa, Barsby & Lithovius et al. 2022 Nat. Biotech.* data. EC = enterochromaffin -like cells. Red lines, median.

To verify expression of the tracer receptors/carriers in SC-islets, we reanalyzed single cell RNA sequencing data of 6-month SC-islet grafts (*2*). *GLP1R* was expressed mostly in the beta- and delta cells as expected (*42, 43, 50*), but also in enterochromaffin -like cells (EC-cells), an endocrine impurity known to be present in SC-islet grafts (*2, 3*) (Fig. 1C, Fig. S1). The amino acid transporter *SLC7A5* (a carrier for [^18^F]FDOPA) was expressed at low levels in all cell types. However, *DDC*, the first and rate limiting enzyme in the metabolism of DOPA was expressed at the highest levels in EC-cells, although expression was seen in all other endocrine cell types (Fig. 1C).

### *KCNJ11^+/+^, KCNJ11*^−/−^ and *KCNJ11*^+/R201H^ SC-islets exhibit expected phenotypes *in vitro*

To allow later examination of the consequences of beta cell functional status on tracer uptake, we sought to engineer SC-islets that are hyper- and hypoactive in their insulin secretion by genome editing the K_ATP_-channel gene *KCNJ11*, critical for triggering insulin secretion (*51*). We edited the *KCNJ11* locus to harbor a homozygous knockout of the gene (*KCNJ11*^−/−^), and separately, introduced a gain-of-function mutation (*KCNJ11*^+/R201H^) (Fig. 2A and Fig S2). These mutations are known to cause CHI and persistent neonatal diabetes, respectively. To validate the three phenotypes, we quantified insulin secretion responses *in vitro* to multiple glucose concentrations and the sulfonylurea glibenclamide (GBC), which acts by closing the K_ATP_-channels. *KCNJ11*^+/+^ increased insulin secretion in response to high glucose, GBC and KCl as mature SC-islets would (*2*). *KCNJ11*^−/−^ SC-islets secreted 3 times more insulin in low glucose compared to *KCNJ11*^+/+^ SC-islets (Fig. 2B) and were unresponsive to GBC (Fig. 2C), replicating our previous model of CHI (*53*). Conversely, the *KCNJ11*^+/R201H^ were unresponsive to glucose, but hyper-responsive to GBC, as expected (*52*) (Fig. 2C). These SC-islets matched our previously thoroughly characterized SC-islets in their cytoarchitecture and proportions of beta and alpha cells. (Fig. 2DE). The *KCNJ11^−/−^* mutation slightly biased differentiation towards beta cells, at the expense of alpha cells, similar to our previous CHI model (*53*) (Fig 2E). The SC-islets thus recapitulated the hyper- and hypoactive phenotypes for later correlation with PET.

**Figure 2.**
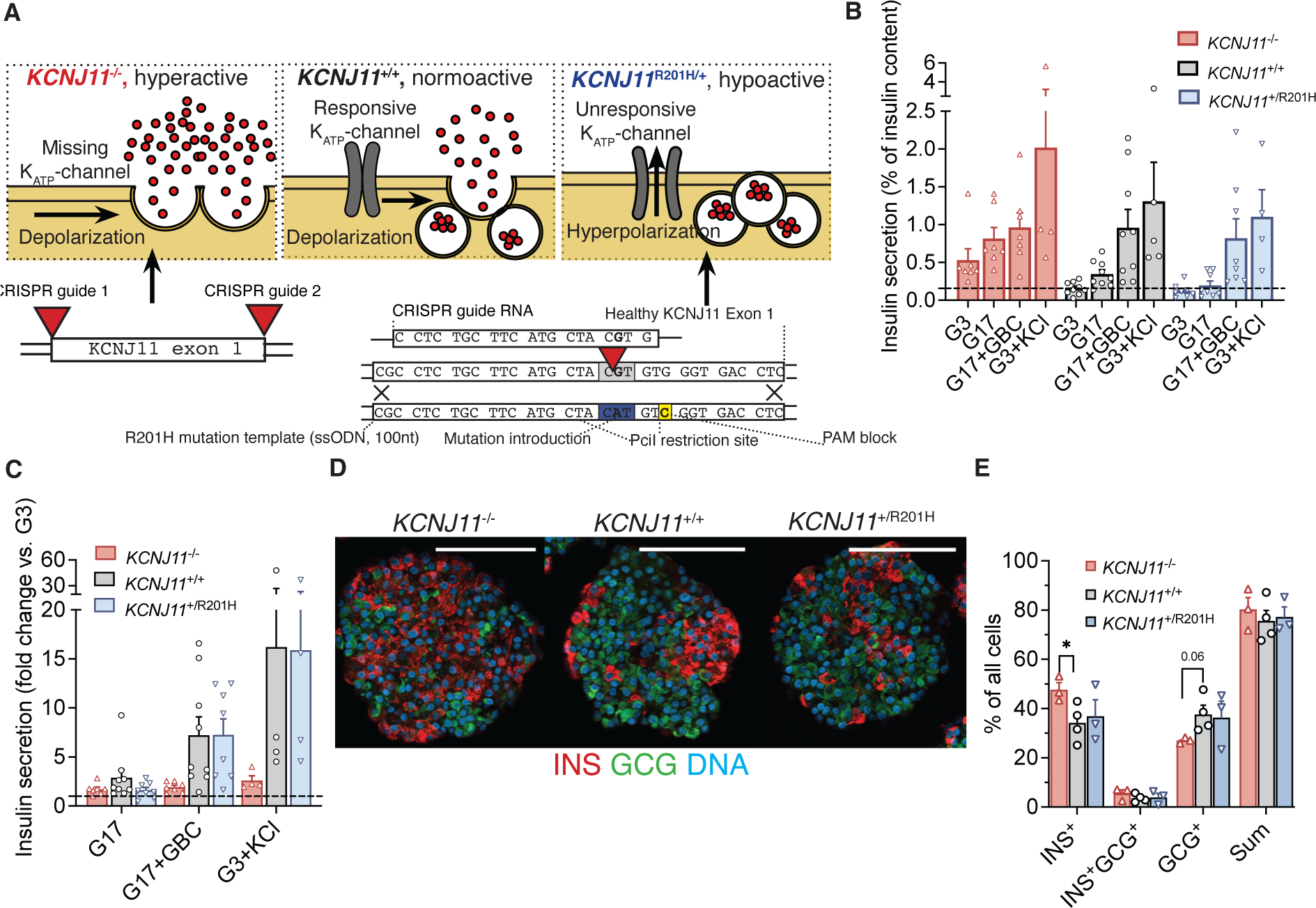
cohort-2 SC-islet phenotypic characterization *in vitro*: **A**) Simplified molecular mechanism and genome editing strategy underlining the hyper-, normo- and hypoactive phenotypes of SC-islet beta cells in cohort-2. *KCNJ11*^−/−^-mutation leads to loss of K_ATP_-channel expression leading to constant depolarization and insulin secretion. *KCNJ11*^R201H/+^-mutation prevents K_ATP_-channel closure under high glucose, leading to inactive beta cells. ssODN = single-stranded oligonucleotide. PAM = protospacer adjacent motif **B**) Percentage of total insulin content secreted in 2.8 mM (G3) or 16.7mM (G17) glucose or G17+ 100 nM glibenclamide (GBC) or G3 + 30 mM KCl. Horizontal line marks mean secretion rate in G3 in *KCNJ11*^+/+^ cells. Tested after 6 weeks of maturation culture **C**) Same as B) normalized against secretion in G3 **D-E**) Immunohistochemisty D) and quantification E) for insulin (INS^+^) beta cells and glucagon (GCG^+^) alpha cells at 6 weeks of maturation culture, scale bars 100 µm. Mean±SEM, One-way ANOVA.

### Engrafted SC-islets develop either into large impure grafts or small pure grafts

After the final 5-month imaging timepoint, the mice were euthanized and the graft-bearing muscles retrieved. The grafts were sometimes visible macroscopically as lightly colored areas (Fig. 3A). We undertook histological analysis for the grafts (n=36), to allow correlation with quantifications from PET. We calculated the volume of the SC-islet grafts from the depth of the grafts as measured by sectioning through the graft-bearing muscle, and from graft area as quantified using the contrast between sparsely nucleated muscle and densely nucleated graft. Total and “cyst-free” graft areas were determined separately (Fig. 3B). The grafts either consisted of SC-islets with interspersed muscle fibers or a fused, heterogenous graft with cysts (Fig. 3B, S3A). SC-islet graft volume was highly variable, ranging from <1 mm^3^ to 150 mm^3^, compared to implanted volume of 1 to 5 mm^3^ (Fig. 3C, S3B). This is likely the result of two stochastic processes: initial cell loss upon engraftment into muscle, a challenging implantation site (*28*); and subsequent growth of the grafted cells. Both are related to maturity of the cells, as more mature islet cells are more susceptible to hypoxia (*29*) and have lower proliferation capacity (*54*).

**Figure 3.**
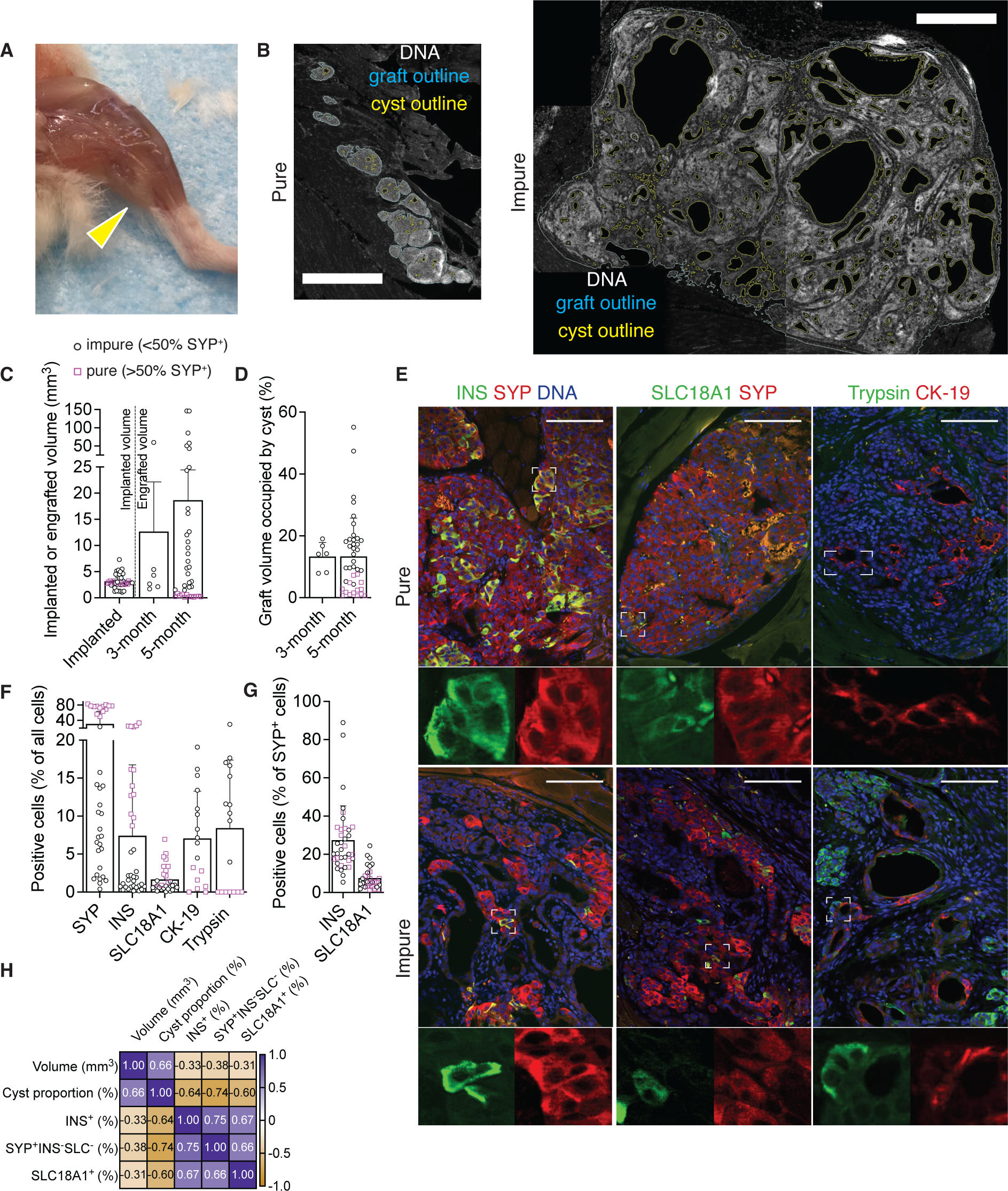
Characterization of the SC-islet grafts: **A**) Skinned mouse hind leg at 5 months post-implantation, arrow indicates graft **B**) Hoechst stained muscle graft sections used for volume determination, examples of pure and impure grafts in the same scale (bar=1000 µm). Displayed as CellProfiler pipeline output images higlighting the total (blue) and cyst-free (yellow) area **C**) Implanted SC-islet volume and histologically determined graft volume. Pink squares = “pure” with >50% synaptophysin^+^ (SYP^+^) endocrine cells out of all graft cells, black circles = “impure” with <50% SYP^+^ **D**) Cyst proportion of the grafts, quantified from images like B). **E**) Immunohistochemistry for SYP, beta cell marker insulin (INS), enterochromaffin cell marker SLC18A1, exocrine marker trypsin and ductal marker cytokeratin-19 (CK-19), scale bar=100 µm, insets 4x magnification. Examples of pure and impure normoactive grafts **F**) Quantification of E) as percentage of all cells in the graft including connective tissue **G**) Percentage of INS^+^ and SLC18A1^+^ cells of SYP^+^ cells **H**) Pearson’s correlation of the composition features

Fluid-filled cysts and ductal mRNA expression are reported to be present in SC-islet derived grafts (*2, 10, 12–15*)). Cyst proportion, quantified here as the percentage of cyst volume of the total volume, was highly variable in the grafts, ranging from around 1% to 50% (Fig. 3D, S3C). The contribution of synaptophysin-positive (SYP^+^) endocrine cells of all graft cells (including connective tissue) separated the grafts in two groups. Many of the grafts had around 80% SYP^+^ cells (“pure” grafts), but the rest had less than 20% SYP^+^ cells (“impure” grafts) (Fig. 3EF, S3D). All the pure grafts were derived from the state-of-the-art Barsby et al. 2022 protocol (*49*), but occasional impure grafts were generated as well.

The putative main target cells of the tracers (namely INS^+^ beta cells and SLC18A1^+^ EC-cells) were found in all grafts. INS^+^ cells represented 10-25% of all cells (including connective tissue cells) in the pure grafts, but less than 5% in the impure grafts (Fig. 3EF, S3D). SLC18A1^+^ cells were present in all grafts at around 1 to 5% of all cells (Fig. 3EF, S3D). Among endocrine cells, INS^+^ cells contributed around 30% and SLC18A1^+^ cells around 10% (Fig. 3G, S3E). Much of the non-endocrine tissue represented connective tissue, but we also analyzed other pancreatic lineages, such as trypsin^+^ exocrine cells and CK-19^+^ ductal cells. No exocrine tissue was detected in the pure grafts, but it was present in the impure grafts (Fig. 3EF, S3D). The cysts were positive for CK-19, indicating the expected pancreatic ductal origin. Even the pure grafts that were devoid of organized cysts had interspersed CK-19^+^ cells representing around 1% of all cells (Fig. 3F, S3D). Large graft volume correlated with high cyst proportion and low numbers of endocrine cells (Fig. 3H). Pure *KCNJ11*^−/−^ grafts had slightly higher beta cell proportion among endocrine cells (Fig. S3F) than *KCNJ11*^+/+^ grafts, similar to what we have described previously (*53*).

In summary, the grafts were highly variable in size, cyst proportion and purity. This allowed graft composition to be explored in the context of tracer uptake, with high clinical relevance regarding the safety and effectiveness of therapeutic SC-islet grafts.

### [^18^F]exendin and [^18^F]FDOPA PET accurately quantify graft volume

As described, we followed the mice for 5 months post-implantation with PET, followed by histological analysis (Fig. 4A). Uptake of both [^18^F]exendin and [^18^F]FDOPA was detected *in vivo* in calf muscles where the SC-islets were implanted (Fig. 4B), while the grafts were undetectable with CT alone. The signal-to-background ratio used to draw the borders of the uptake volume (Fig. 4C), was slightly higher for [^18^F]exendin compared to [^18^F]DOPA (Fig. 4D). Some grafts were already detectable at 1-month post-implantation but the detection rate improved over time to around 90% for exendin and 70% for DOPA at the final 5-month imaging timepoint (Fig. 4E). The volumetric resolution of the PET scanner is around 1 mm^3^, so we used that as a cutoff when calculating the detection rate in relation to graft size. All but one graft >1mm^3^ were detected with [^18^F]exendin (96% detection rate) with [^18^F]FDOPA also performing well (85% detection rate) (Fig. 4F). The grafts <1mm^3^ were better detected with [^18^F]exendin (69 vs. 44%), suggesting that uptake of [^18^F]exendin in these small but pure grafts is so high that the signal spillover overcomes the theoretical detection limit (Fig. 4F).

**Figure 4.**
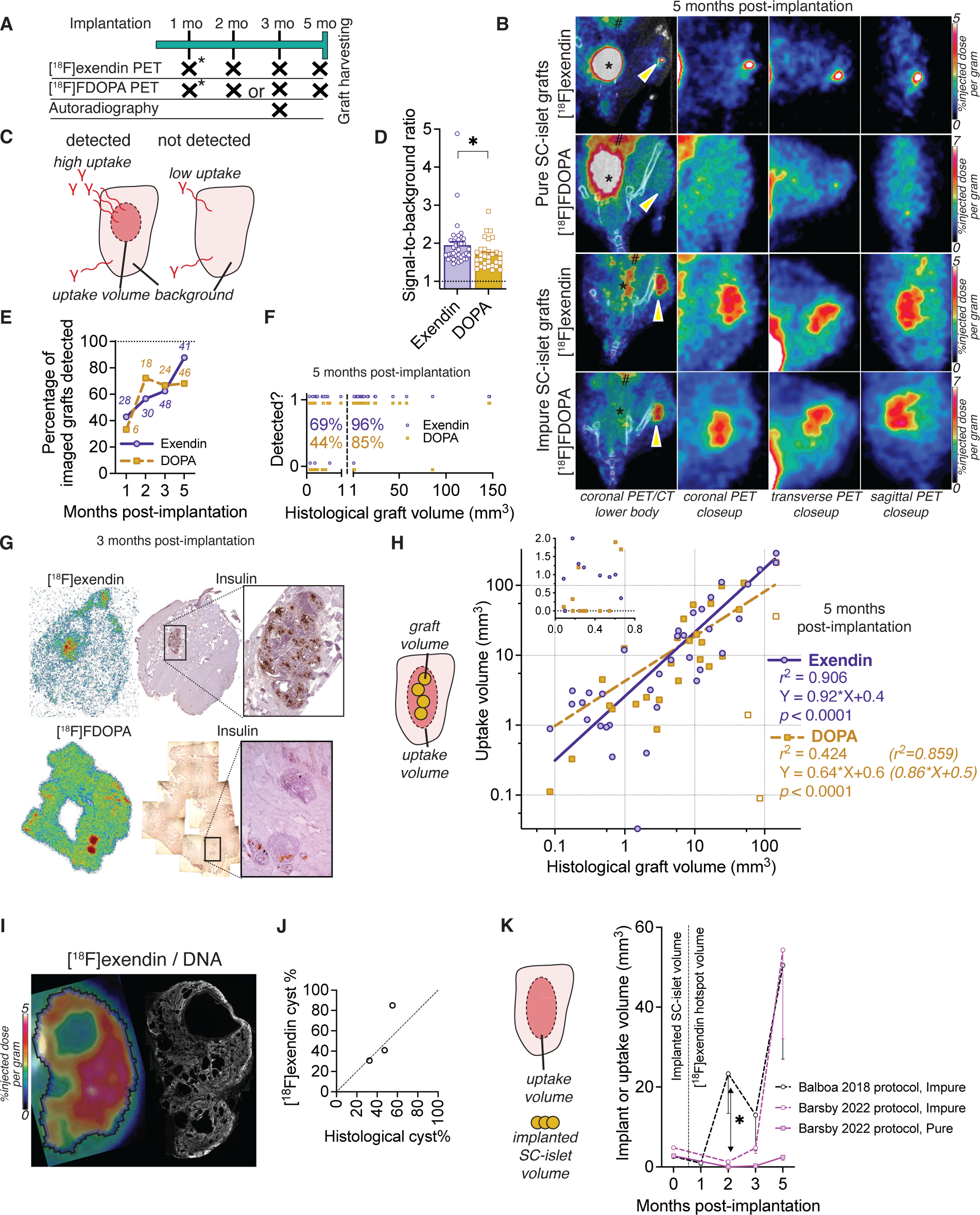
follow-up and quantification of SC-islet graft volume with PET: **A)** Imaging follow-up, crosses indicate imaging or ex vivo autoradiography timepoint.*cohort-1 only **B**) Views of PET/CT and PET images (summed 30-min scans), arrows indicate candidate uptake volume * bladder, # the kidneys **C)** Illustration of graft detection based on gamma emission in the muscle (γ) corresponding to tracer uptake **D**) Ratio of tracer uptake concentration inside the uptake volume and in the surrounding muscle at 5 months post-implantation, Welch’s t-test **E-F**) Tracer detection performance, E) Per timepoint (*N* on graph) F) Per histological volume (1=detected, 0=not detected). **G)** *Ex vivo* autoradiography of muscle sections taken immediately after imaging with [^18^F]exendin or [^18^F]FDOPA and insulin immu-nohistochemistry from adjacent sections **H)** Linear regression between [^18^F]exendin and [^18^F]FDOPA uptake volume and the histologically determined graft volume (including cysts). Inset highlights low values. Values in brackets are for [^18^F]FDOPA when three of the most cystic grafts are excluded (white-filled squares) **I**) Example frame of [^18^F]exendin-PET and Hoechst stained section of the same graft **J**) Cyst proportion determined with histol-ogy or PET (by drawing full and cyst-free uptake volume) **K**) Graft size follow-up with [^18^F]exendin PET, grouped by purity and SC-islet differentiation protocol, with grafts having >50% of all cells synaptophysin^+^ considered pure. Unpaired Welch’s t-test. *N*: *Balboa* impure=17, *Barsby* impure=11, *Barsby* pure=13

Both tracers reached a steady-state concentration within 5 minutes after injection, which remained on a high level for the 30-minute scan, suggesting the uptake corresponded to tracer-targeted SC-islet tissue (Fig. S4B). For further evidence of tracer graft specificity, we sacrificed a subset of mice immediately after PET-imaging for *ex vivo* autoradiography. The autoradiography sections showed uptake of both [^18^F]exendin and [^18^F]FDOPA, corresponding to locations staining positive for insulin in adjacent sections (Fig. 4G), verifying uptake in the graft-bearing muscle is specific to the SC-islet graft.

For PET to be a reliable method for graft monitoring, graft volume determined with PET should correspond to actual graft volume. We thus correlated the uptake volume determined by both [^18^F]exendin and [^18^F]FDOPA with the histologically determined graft volume. The uptake volume quantified with [^18^F]exendin correlated very strongly (*r*^2^=0.91) with actual graft volume (Fig. 4H), indicating high accuracy for graft volume quantification in grafts of vastly different sizes and purities. [^18^F]FDOPA uptake volume also strongly correlated with actual graft volume when analyzing all but the most cystic grafts (*r*^2^=0.86). However, the precise outlines of three highly cystic grafts were difficult to determine with [^18^F]FDOPA, reducing the strength of correlation when included (*r*^2^=0.43) (Fig. 4H). Their outline and the largest individual cysts inside them were readily visualized with [^18^F]exendin (Fig. 4I, S4CD). By including or excluding the cysts in the uptake volume, we could noninvasively determine the cyst proportion, which closely resembled the one determined with *ex vivo* histology of these three grafts (Fig. 4J).

Finally, as we verified the graft volume determined by PET did correspond to actual graft volume, we analyzed the growth patterns of the scanned SC-islet grafts over the 5-month follow-up. The grafts displayed consistent patterns of growth or stability in the uptake volume, with tracer uptake concentration also remaining relatively constant during follow-up (Fig. S4EF). Interestingly, after grouping the grafts based on final endocrine purity, the impure grafts derived with the Balboa 2018 protocol showed unwanted growth at 2 months post-implantation, whereas growth of the occasional impure grafts derived from the Barsby 2022 protocol showed expansion later at the 5-month timepoint (Fig. 4K). The volumes of grafts that remained pure at 5 months (all derived with Barsby 2022 protocol) were stable in longitudinal imaging (Fig. 4K). These results show that the starting SC-islet material influences the temporal growth pattern of SC-islet grafts, and that these patterns can be detected using PET.

Taken together, these data demonstrate the validity of [^18^F]exendin and [^18^F]DOPA PET for longitudinal monitoring and quantifying the actual SC-islet graft volume noninvasively across a wide distribution of graft volumes and purities.

### [^18^F]exendin uptake estimates graft beta cell proportion and is unaffected by beta cell functional state

We then investigated which aspects of graft composition and function determine uptake of the tracers; or conversely, if tracer uptake can be used to inform about compositional or functional aspects noninvasively. As described above, the pure grafts were very small (<1 mm^3^ in volume) but still visible due to high uptake of the tracers spilling over to the surrounding muscle. In this situation, much of the uptake volume consists of the muscle, diluting the measured graft uptake concentration. However, with background uptake measurements and histologically determined graft volume, we could calculate graft volume specific uptake concentration, correcting for the dilution (Fig. 5A, detailed in the methods).

**Figure 5.**
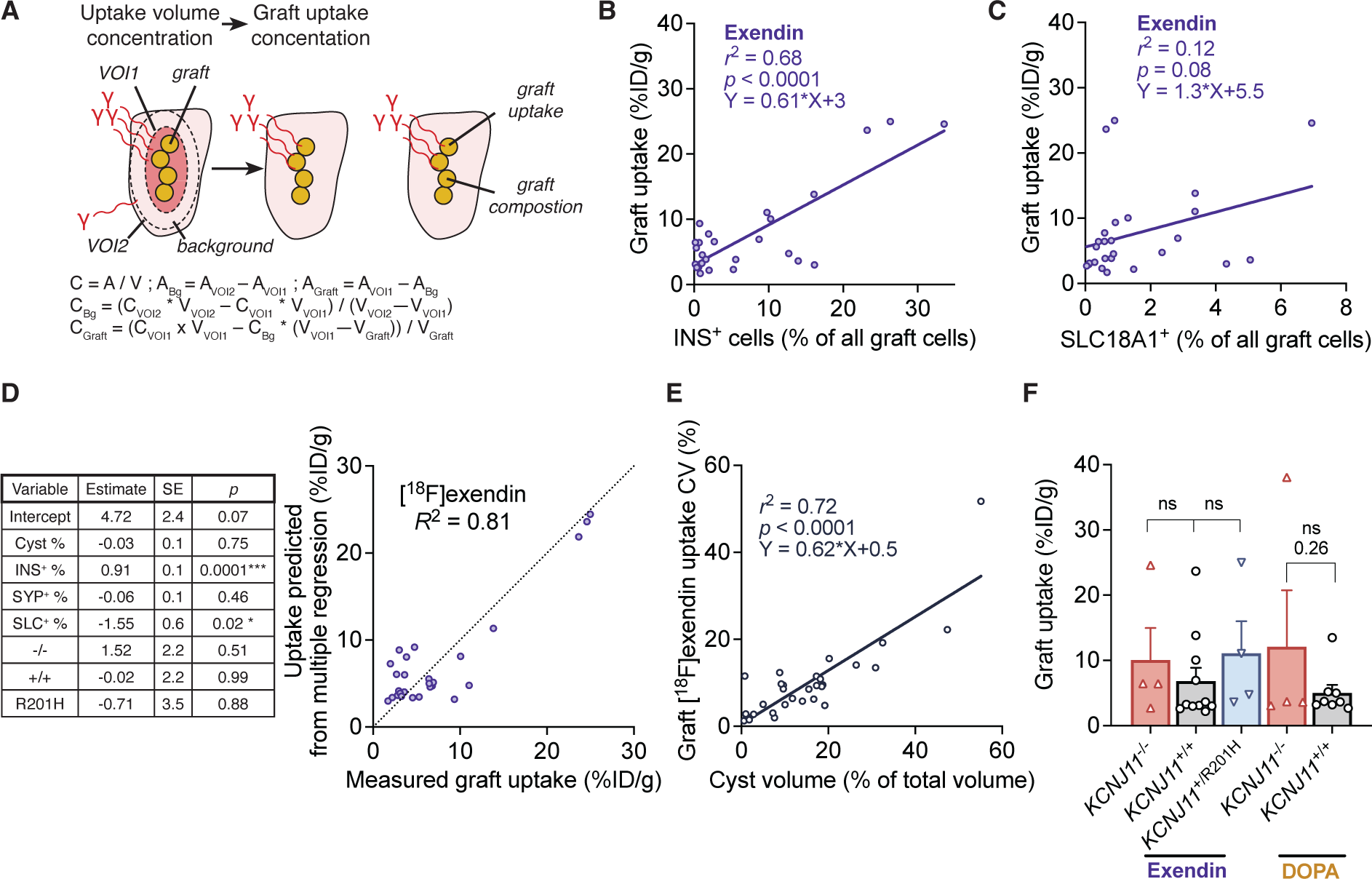
Correlation of graft composition and genotype with tracer uptake: **A**) Schematic and associated formulas of volume dilution correction to determine graft uptake concentration. C = tracer uptake concentration (%ID/g: percentage of injected tracer dose per gram of hotspot), A=radioactivity (becquerel), V = volume (mm^3^), Bg = background, VOI1= uptake volume, VOI2=additional VOI encompassing the whole muscle **B-C**) Linear regression of graft exendin uptake and graft beta cell B) and EC cell C) proportion **D**) Multiple linear regression of tracer uptake from composition, in the form: Y (Predicted uptake) = β_intercept_+β_Cyst_*X_Cyst_+β_insulin_*X_insulin_… (X area histological measurements from individual grafts). Table: Model estimates of β, SYP^+^ = SYP^+^INS^-^SLC18A1^-^, SLC^+^ = SLC18A1^+^, −/−, +/+ and R201H refer to *KCNJ11* genotype. Graph: Correspondence of values from model to measured uptake and model goodness-of-fit **E**) Linear regression of graft [^18^F] exendin uptake variability and cyst proportion. CV (coefficient of variation) = Uptake SD / Graft uptake mean *100 **F**) Graft [^18^F]exendin and [^18^F] FDOPA uptake in cohort-2 grafts of different genotypes. A single *KCNJ11*^+/R201H^ graft was detected with DOPA and is not plotted.

To explore what aspects of the graft composition determine [^18^F]exendin and [^18^F]FDOPA uptake, we correlated graft uptake concentration with histologically determined cyst proportion and cell type percentages from the same graft. Simple Pearson’s correlation of tracer uptake and composition parameters showed negative correlation with cyst proportion and positive correlation with beta and EC cell proportions (Fig. S5A). Analyzed individually with linear regression, graft INS^+^ beta cell percentage positively correlated with both [^18^F]exendin (*r*^2^=0.68) and [^18^F]FDOPA (*r*^2^=0.52) uptake (Fig. 5B, S5B).

The number of SLC18A1^+^ EC cells was associated with higher uptake of [^18^F]FDOPA only (*r*^2^=0.48) (Fig. 5C, S5C). Graft uptake of [^18^F]exendin and [^18^F]FDOPA did not correlate with the percentage of SYP^+^INS^-^SLC18A1^-^ cells (i.e. alpha and delta cells) (Fig. S5D). As multiple factors contributed to uptake of both tracers, we analyzed them simultaneously using a multiple regression model, in the form of *Uptake ∼ Intercept + Cyst% + INS^+^% + SYP^+^INS^-^SL18A1^-^% + SLC18A1^+^% + Genotype*. This regression model had a better fit with graft [^18^F]exendin uptake (*R*^2^=0.81) than any of the parameters alone (Fig. 5D). Percentage of INS^+^ beta cells was the most important determinant for [^18^F]exendin uptake in the model (Fig. 5D). Analyses on [^18^F]FDOPA relied on fewer observations as most of the pure and small grafts were undetected, and we state conclusions on its uptake determinants with less confidence. These data show that the real-world determinants of tracer uptake correspond well to the pattern predicted based on tracer target expression in single-cell RNA sequencing (Fig. 1C). Our observations also demonstrate the ability of PET to inform about multiple aspects of SC-islet graft composition, e.g. their beta cell percentage.

All the impure grafts had cysts, with three having so large they could be clearly distinguished with [^18^F]exendin (Fig. 4I). In addition to this, uptake volume in the other impure grafts had highly variable uptake concentration pattern inside each graft, which could be translated into the coefficient of variation of tracer uptake. This [^18^F]exendin uptake variation correlated with cyst proportion (Fig. 5E) (*r*^2^=0.72, using graft uptake for the calculations), suggesting cyst proportion could be estimated noninvasively even when individual cysts were too small to be visualized directly.

We then explored how the functional status of engrafted beta cells influences tracer uptake. The cohort-2 mice carried grafts with a hyperactive *KCNJ11*^−/−^ or a hypoactive *KCNJ11*^+/R201H^ graft in one leg and a normoactive *KCNJ11*^+/+^ graft in the other. When comparing graft tracer uptake concentration in these functionally different grafts, no significant differences were found (Fig. 5F). The multiple regression model also pointed towards a low influence of genotype in determining tracer uptake (Fig. 5D). The graft genotype did not affect the uptake dynamics of either [^18^F]exendin or [^18^F]FDOPA (Fig. S5EF). This indicates that the tracers can be confidently used in SC-islet grafts exhibiting various states of beta cell functionality as even the phenotypic extremes used here did not bias the tracer uptake. Taken together, the composition of the grafts, namely the contribution of beta and EC-cells, determine the level of uptake in the grafts, rather than the functional status of the beta cells.

### Graft volume and beta cell mass cannot be quantified with C-peptide measurements

Finally, to contextualize the PET-imaging results, we explored the ability of C-peptide measurements taken under fasted, glucose-stimulated and hypoglycemic conditions to quantify graft volume and beta cell mass. The imaged mice carried two grafts, and so were not suited for correlating against the circulating C-peptide originating from both grafts. The sum of grafts’ beta cell mass had no correlation with fasting C-peptide (Fig. S6A). To explore this further in the most controlled way, we implanted an additional cohort of mice (n=14) with SC-islets under the kidney capsule in three size categories (1, 3 or 6.3 mm^3^, 1 mm^3^ = 566 islet equivalents ≈ 150 SC-islets), all coming from one batch of SC-islets derived with the Barsby et al. 2022 protocol.

The C-peptide levels from the kidney grafts were higher overall than in the intramuscular grafts (Fig. S6B). In 5-month follow-up, the kidney grafts increased their fasting C-peptide secretion and lowered the recipients’ fasting blood glucose level to human levels as expected (*2*), with minor differences between the size groups (Fig. 6AB). Remarkably, even the smallest grafts caused human level fasting blood glucose and high C-peptide levels, albeit after a longer lag period. We then performed glucose- and insulin tolerance tests at 5- and 5.5-months post-implantation. In the glucose tolerance test (Fig. 6CD), the mice with the lowest implanted dose (1 mm^3^) did not show glucose-dependent C-peptide secretion, while the mice carrying larger grafts showed 2 – 3 fold responses (Fig. 6D, inset). The 1 mm^3^ dose was thus subtherapeutic, similar to what was found in a study comparing 0.75 vs. 2 and 5 million implanted SC-islet cells (*28*). In the insulin tolerance test, all grafts efficiently shut down their insulin secretion when the mice became hypoglycemic (Fig. 6E), with around 90% reduction from baseline (Fig. 6F, inset). These data indicate the SC-islet grafts are functional *in vivo* displaying glucose regulated stimulation and shutdown of insulin secretion.

**Figure 6.**
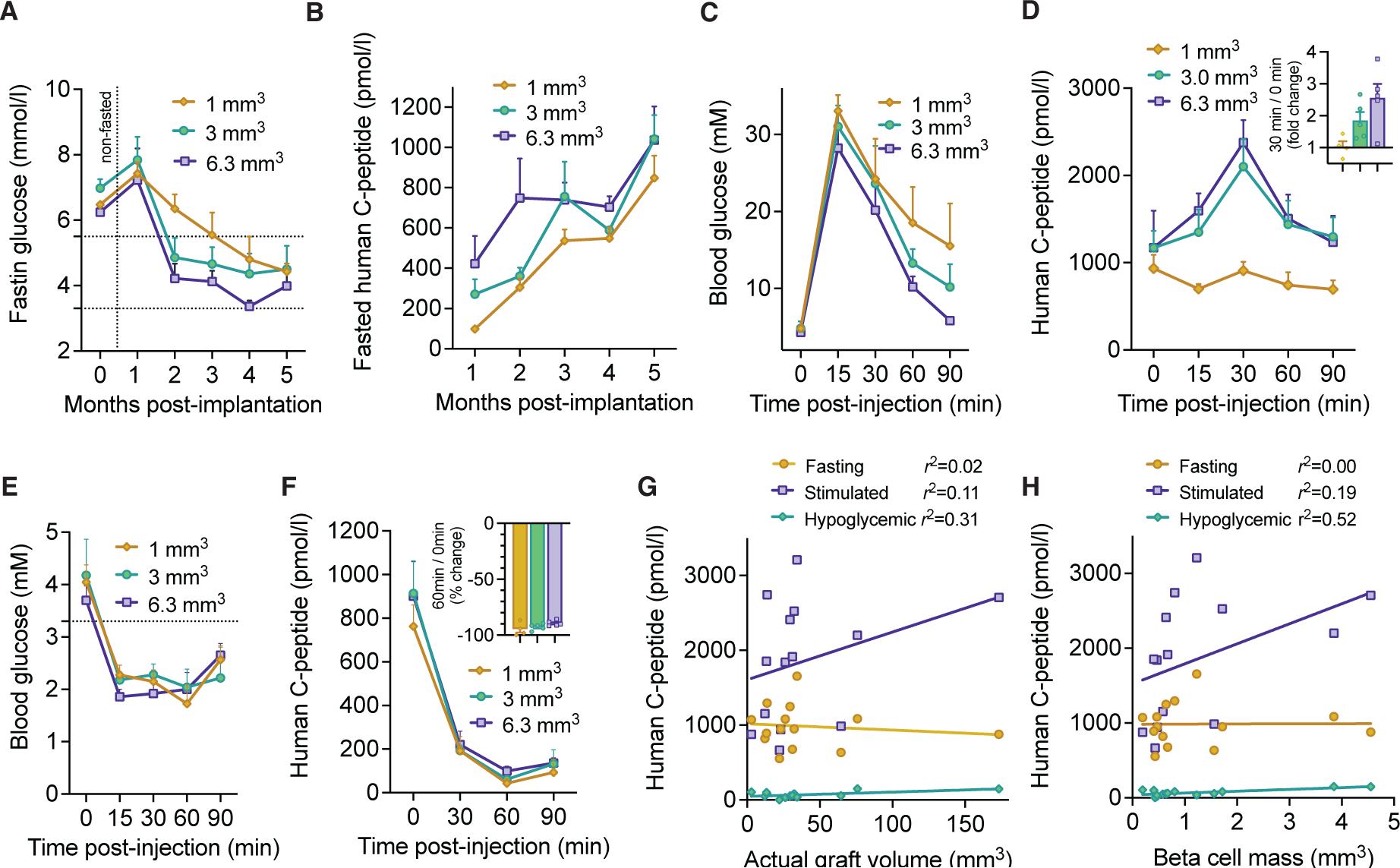
relationship between C-peptide and graft volume: **A-B**) Follow-up of fasting glucose A) and fasting C-peptide B) in the kidney cohort, grouped by total implanted SC-islet volume. Normal human fasting blood glucose level marked. **C-D**) Glucose C) and C-peptide D) following intraperitoneal injection of glucose (3 mg/g) at 5-months post-implantation **E-F**) Glucose E) and C-peptide F) following intraperitoneal injection of insulin analogue (Actrapid) (0.75 mIU/g) at 5.5-months post-implantation **G-H**) Linear regression of C-peptide parameters and total volume of the kidney graft G), or the cyst-free, beta cell fraction corrected graft volume H). The C-peptide parameters were: C-peptide at the fasted state (mean of two days), or at 30 minutes after the glucose injection, or at 60 minutes after the insulin injection

Following the insulin tolerance test, graft-bearing kidneys were harvested, and actual graft volume, cyst proportion and beta cell percentage was determined histologically, as in the intramuscular grafts (Fig. S6C). Importantly, we then correlated the actual graft volume against various C-peptide measurements from the same mice: namely, the final fasting C-peptide level, peak C-peptide in the glucose tolerance test and nadir C-peptide in the insulin tolerance test. None of these measures correlated with actual graft volume, which included cysts and non-beta cells that do not secrete C-peptide (Fig. 6G). We then determined the graft beta cell mass, by subtracting their cyst volume and multiplying the remainder with the histologically determined beta cell percentage. Beta cell mass weakly correlated with the remaining C-peptide secretion under hypoglycemia (*r*^2^=0.52), but not with fasting- or stimulated C-peptide (Fig. 6H). These results indicate that C-peptide measurements cannot accurately quantify the total graft volume, unlike PET imaging.

## DISCUSSION

In this study, we show that PET-imaging with ^18^F-labeled exendin and DOPA enables detection and follow-up of human stem cell derived islet grafts in mice. By measuring the volume of tracer uptake, we were able to quantify the actual volume of the grafts, be they pure or impure. Additionally, the characteristics of tracer uptake yielded information about the fraction of beta cells and cysts, enabling noninvasive study of SC-islet graft size and composition.

We imaged the SC-islet grafts with both [^18^F]exendin and [^18^F]FDOPA, providing unique perspective on their strengths and weaknesses as tracers for SC-islet imaging (Table 1). The tracers performed similarly overall with a strong ability to quantify actual graft volume, although minor differences were found. [^18^F]exendin was more sensitive than [^18^F]FDOPA in detecting the grafts, especially the small and pure ones. This heightened sensitivity was also reflected in the ability of [^18^F]exendin to detect large cysts inside the grafts, as well as detecting the presence of smaller cysts by the increased [^18^F]exendin uptake variability. The uptake of [^18^F]exendin was predominantly determined by the proportion of beta cells in the graft, while [^18^F]FDOPA uptake appeared to be determined by the proportion of beta cells and the less numerous EC-cells. In practice then, a patient imaged with either tracer could have the volume of their graft quantified based on the uptake volume. If a large uptake concentration would be measured, a higher number of beta cells could be inferred to be present in the graft, at least when using [^18^F]exendin. [^18^F]FDOPA has an important practical advantage being readily available in PET-centers worldwide for use in central nervous system and focal CHI imaging (*45*). PET with ^68^Ga- or ^18^F-labeled exendin on the other hand is limited to a handful of dedicated hospitals in Europe and Japan. The uptake of neither tracer was biased by the extremes of beta cell insulin secretion functionality. This is reassuring, as some degree of functional variability is bound to be seen in ongoing and future clinical SC-islet grafts.

**Table 1.** comparison between [^18^F]F-DBCO-exendin-4 and [^18^F]FDOPA.

| Parameter | [$^{18}\text{F}$]exendin | [$^{18}\text{F}$]FDOPA |
| --- | --- | --- |
| Detection of grafts <1 mm<sup>3</sup> | ++ | + |
| Detection of grafts >1 mm<sup>3</sup> | +++ | ++ |
| **Quantification of graft volume** | **+++** | **++** |
| Assessment of cyst proportion | + | 0 |
| Influence of beta cell percentage on tracer uptake | ++ | + (?) |
| Influence of enterochromaffin cell percentage on tracer uptake | 0 | + (?) |
| Influence of beta cell functional state on tracer uptake | 0 | 0 |
| **Clinical availability** | **0** | **++** |

*Authors' assessment on performance in SC-islet graft context: +++ strong ability/effect, ++ moderate ability/effect, + marginal ability/effect, 0 no ability/effect, (?) lower level of confidence*

The reason for increased [^18^F]FDOPA uptake in focal CHI is currently unknown. Two explanations are theorized: firstly, the hyperfunctionality of the lesion beta cells increasing their demand for metabolites like DOPA; or secondly, local increase the number of beta cells. Based on our data of CHI grafts having similar [^18^F]FDOPA uptake as healthy and diabetic grafts, the reason for increased [^18^F]FDOPA uptake in the lesion is most likely the locally large number of beta cells. Pointing to the same conclusion, Boss and colleagues found that focal CHI lesions can be located with exendin PET better than with DOPA PET (*55*). According to our data, this performance benefit would be due to the improved ability of exendin in quantifying graft volume, not due to differential uptake in hyperactive grafts. However, in cases of extreme hyperglycemia, GLP1-receptor expression does increase, which has been shown to influence exendin tracer uptake at least in the native pancreas (*56*).

The overall purity differences in the SC-islets strengthened the conclusions of our imaging data but pose a challenge for SC-islet translation into clinical use. Most of the impure grafts in our study were derived from SC-islets differentiated with previous generation protocol. Nevertheless, a batch of SC-islets derived with our state-of-the art protocol also generated cystic, impure grafts. The precise origin of the unwanted growth remains an important question in the field, with reports pointing to lingering pancreatic progenitor cells capable of ductal and exocrine differentiation and non-pancreatic contamination, resulting in duodenal- and liver-like cells (*14, 15*). Rigorous pre-implantation quality control for the contaminating cell populations is needed, as functional maturity does not necessarily equate a risk-free graft. The imaging methods described here could be useful tools to supplement pre- implantation quality control informing decisions to intervene, if the graft becomes too large or too impure.

C-peptide level has been reported to be independent of transplanted primary human islet mass in mouse (*34*) and human recipients (*35*). Similarly, in our study the therapeutic doses produced similar C-peptide levels in mice and showed the inability of C-peptide to measure overall graft volume. Our study also investigated the connection of C-peptide with engrafted beta cell mass. Both fasted and glucose stimulated C-peptide measurements were poor predictors of beta cell mass. Upon induced hypoglycemia the grafts efficiently shut down insulin secretion by 90% from fasted levels, indicating effective glucose sensing. Despite this, we could still detect low levels of inappropriate insulin release. Interestingly, this insulin leak was a weak indicator of beta cell mass.

A clear advantage of PET is its direct translatability to clinical use. Importantly, the larger human scale would likely greatly improve the ability of this technique to quantify relevant changes both in graft volume and composition, since the human scanners operate on approximately three-fold lower resolution, but the graft size would need to be a thousand-fold larger for therapeutic effectiveness. Indeed, assuming the islet dose of around 15,900 islet equivalents per kg as was the mean dose in patients achieving sustained graft survival in islet transplantation (*1*), the total implanted SC-islet volume would be almost 2000 mm^3^ for a 70 kg person (1 islet equivalent = 0.00177 mm^3^). In our study, the graft volume in many of the grafts was at the lower limit of the PET scanner’s spatial resolution (of around 1 mm^3^), but the grafts could still be detected. The larger graft volumes would result in lower dilution in uptake concentration likely making quantification of graft composition more accurate than in this study. ^18^F-labeled exendin has been shown to have low enough radiation dose where even repeated scans could be justified (*57*). Altogether, it should be much more feasible to conduct SC-islet graft PET-imaging in human graft recipients than what we achieved in mice.

One obstacle to translating these findings to human use is the use of intraportal transplantation as the standard strategy, also employed in one of the ongoing clinical SC-islet implantation trials (*NCT04786262*). This is a challenging site to image as the islets engraft next to the large blood vessels supplying the liver, leading to background signal from the tracer in circulation. However, in a promising outcome for applying PET for intrahepatic SC-islet grafts, a recent study showed that primary islet grafts can be detected in the liver (*39*) with a ^68^Ga-exendin tracer. Additionally, while our method was highly applicable for long-term monitoring it was limited in mapping the early post-implantation events, as many of the grafts were not yet detected at the 1-month timepoint. These events can well be explored in a preclinical setting, where bioluminescent reporter systems can be used.

As previously detailed, strengths of this study include its setup of comparing two different PET-tracers; inclusion and quantification of grafts with different compositions allowing identification of tracer uptake determinants; validation of tracers in SC-islets with hyper- and hypofunctional beta cells; as well as the comparison to C-peptide based measurements. In conclusion, we propose PET imaging utilizing the tracers [^18^F]exendin and/or [^18^F]FDOPA as a method for longitudinal monitoring of SC-islet grafts. This would allow accurate quantification of the graft volume and estimation of aspects of their composition, such as cyst proportion and beta cell contribution. PET-imaging could help improve the safety and efficacy of SC-islet grafts as an emerging cell replacement therapy for insulin deficient diabetes.

## MATERIALS AND METHODS

### Study design

Initial objectives were: 1) Can SC-islet graft size be quantified with PET? 2) Does SC-islet graft functional state affect PET-tracer uptake? Additional objectives were formulated based on preliminary results: 3) Does SC-islet graft composition influence tracer uptake? 4) Do C-peptide measurements quantify graft size and beta cell mass? This exploratory study had no predetermined sample size.

The study was designed to be conducted in two cohorts of mice implanted with different types of SC-islet grafts. Each cohort comprised of several batches of around 6 mice, limited by the number that could be imaged within tracer half-life. Cohort-1, comprising of 17 mice, was designed to explore the relationship between graft size in PET and its actual size. Mutation-corrected induced pluripotent cell line HEL113.5(*53*) derived SC-islets were used. They were differentiated with the Balboa et al. 2018 protocol, modified to use Aggrewell 400 plates (Stem cell technologies) for aggregation (*48*). Cohort-2, comprising of 18 mice, was designed to explore the relationship between graft functional state and its PET tracer uptake. SC-islets were differentiated from the H1 embryonic stem cell line (WiCell) and its gene-edited counterparts with Barsby et al. 2022 protocol, which yields SC-islets achieving functional maturity *in vitro* (*2, 49*). The kidney cohort was studied as a single batch of 14 mice.

No outliers were excluded. PET data was not available for the non-detected grafts and are missing from further analyses, except on Fig. 4H, where they are plotted with PET volume =0. All replicates are biological, from different batches of SC-islets, mice, or SC-islet grafts. Mice were randomly allocated to have different types of implants. Only female mice were used, and littermate controls preferred. PET imaging was conducted blinded. PET analysis was not blinded. Histological analyses were conducted with an automated, blinded-by-design pipeline.

### Genome editing of the *KCNJ11* locus

The H1 line was separately genome-edited to generate *KCNJ11* knockout and *KCNJ11* R201H-knock-in. Genome editing was performed using ribonucleoprotein (RNP) CRISPR-Cas9 system (Integrated DNA Technologies), with guides and a mutation template designed using Benchling (Biology Software, 2017). Two million cells were electroporated with 10 µg Alt-R™ S.p. Cas9 Nuclease V3 combined with gRNA and 4 µg of the 100-b ssODN mutation template (Integrated DNA Technologies), using Neon Transfection system (Thermo Fisher; 1100 V; 20 ms; two pulses). To knockout *KCNJ11*, we targeted a 1.27 kb deletion spanning the protein-coding exon using 2 gRNAs (KO gRNA1: AGGCCCTAGGCCACGTCCGA and KO gRNA2: GTGTGTACACACGGACCATG). The deletion was confirmed with PCR using the following primers, *KCNJ11* KO Fw: CTCAGCCTCCCAACGTACTG and *KCNJ11* KO Rv: AGAGTGTGGCTGGTCAATCG. For knocking in the *KCNJ11*-R201H gain-of-function mutation, we used KI gRNA: CCTCTGCTTCATGCTACGTG and a mutation template harboring a silent mutation to create a PciI restriction site facilitating screening for edited clones. Monoclonal cells were isolated using limiting dilution, expanded, and screened using PCR and Sanger sequencing, with the following primers used for screening: *KCNJ11* KI Fw: TCATCGTGCAGAACATCGTG and *KCNJ11* KI Rv: ATCTCATCGGCCAGGTAGGA. One homozygous knockout and two heterozygous R201H knock-in clones were produced. The top three CRISPR off-target hits for each gRNA were sequenced demonstrating no insertions or deletions. G-band karyotyping was performed at Ambar Lab, Barcelona, Spain.

### SC-islet implant volume determination

Initial implant preparate volume was determined by hand-picking the SC-islets for each graft on a separate well and taking an image of each well with a stereomicroscope. Each image was then analyzed with a custom CellProfiler pipeline (version 3.0 used)(*58*), which yielded the number of SC-islets in the image as well as their average diameter in millimeters, from which their total volume could be calculated assuming sphericity. A predetermined SC-islet volume was then aspirated with a precision syringe (Hamilton, #81301) into silicone tubing, then compacted into a pellet inside via centrifugation in 100g for 2 minutes and kept on ice until implantation. In cohort-1, four sizes of implants were prepared (volumes in mm^3^): around 1.3; 2.5; 3.8 or 5.0, with the left calf implanted with a smaller graft than the right. In cohort-2, either around 3 or 5 mm^3^ of SC-islets were implanted in each leg. The left leg was implanted with *KCNJ11*^−/−^ or *KCNJ11*^+/R201H^ SC-islets and the right leg with *KCNJ11*^+/+^ SC-islets.

### SC-islet implantation

Immunocompromised female NSG mice were anesthetized with isoflurane (5% induction, 2% maintenance) and both their calves shaved. The tubing containing the SC-islet pellet was connected to another precision syringe and the other end to a 0.6 mm needle (Sarstedt 85.923). The needle was inserted into the distal, posterolateral calf parallel to its long axis and pushed 4-5 mm inside the *musculus gastrocnemius*. The SC-islets were implanted while retracting the needle by 1 mm. After implantation, if any SC-islets remained inside the opaque needle, their volume was quantified and subtracted from implanted volume. Kidney implantations were conducted as previously reported (*2*).

### Animal husbandry and C-peptide measurements

After implantation the mice were first housed for two weeks in Helsinki and then transported to the University of Turku animal facility for 5-month PET imaging follow-up. The kidney cohort used for C-peptide studies was kept in the University of Helsinki animal facility. Both were housed in Scantiner or IVC system, fed irradiated standard chow, and kept on a 12h dark/12h light cycle. C-peptide measurements were taken from *vena saphena magna* or via cardiac puncture. Glucose- and insulin tolerance tests were conducted in the afternoon after a 5-hour fast by injecting 3 mg/g of glucose or 0.75 mIU/g of insulin intraperitoneally, with blood sampling intervals as indicated in the figures. The C-peptide concentration was analyzed with a human specific ultrasensitive ELISA kit (Mercodia, 10-1141-01), as per manufacturer instructions. The animal care and experiments were approved by the Finnish national animal experiment board (ESAVI/14852/2018, ESAVI/9734/2021, ESAVI/12143/2019 and ESAVI/16273/2022).

### Radiotracer synthesis

Radiotracers were synthetized at the radiochemistry laboratory in Turku PET center, University of Turku. [^18^F]FDOPA was synthesized via electrophilic ^18^F-fluorination as previously described (*59*). [^18^F]F-DBCO-exendin-4 (Nle^14^,Cys^40^(Mal-dPEG4-DBCO-N_3_-PEG34-ethyl[^18^F]fluoro-exendin-4) was synthesized in two steps using a prosthetic group strategy (*44*). In the first step the prosthetic group, azido-PEG4-[^18^F]fluoride was synthesized from tosylate precursor and K_222_/[^18^F]/K^+^-complex via nucleophilic fluorination. In the next step, [^18^F]F-DBCO-exendin-4 was synthetized from cyclooctyne-derivatized exendin-4 and azido-PEG34-ethyl[^18^F]fluoride via strain-promoted azide-alkyne cycloaddition. [^18^F]F-DBCO-exendin-4 was purified via semipreparative HPLC and formulated in 10% ethanol in 0.1M PBS containing 0.02% ascorbic acid. Molar activity of [^18^F]F-DBCO-exendin-4 was 40 - 100 GBq/µmol and [^18^F]FDOPA > 3 GBq/µmol at the end of synthesis.

### PET/CT imaging

Mice were fasted for 4h prior to imaging to standardize blood glucose levels. One or two mice were scanned at a time, with small-animal PET/CT (X- and β-cubes, Molecubes NV, Belgium) after intravenous administration of approx. 2 - 3 MBq of either [^18^F]FDOPA or [^18^F]exendin under isoflurane anesthesia. A 10-min CT for attenuation correction and anatomical reference preceded PET. Dynamic PET data were collected for 30 minutes, divided into 25 time frames (12×10s, 6×30s, 5×60s, 2×600s), in list mode and reconstructed with an OSEM3D algorithm. The radioactivity uptake was corrected for radionuclide decay and expressed as percentage of injected dose per gram tissue (% ID/g). The scanner has a 13-cm axial and a 7.2-cm transaxial field of view generating 192 transaxial slices with voxel dimensions of 0.4 x 0.4 x 0.4 mm. The PMOD software (v. 4.0) was used to convert reconstructed DICOM images produced by the β-cube into a Inveon Research Workplace (v. 4.2, Siemens Medical solutions) compatible DICOM format, used for data analyses.

### PET data analysis

The image frames were summed and scaled from 0 to 5 %ID/g for [^18^F]exendin and 0 to 7 %ID/g for [^18^F]FDOPA and volumes of interest (VOI1) were drawn when hotspots were detected in the calf muscle. VOI inclusion was determined by contrast between background uptake and high tracer uptake areas as displayed in results. Areas inside the VOI displaying lower uptake were included for all grafts. Three grafts were additionally quantified by excluding areas inside with low uptake.

Larger VOIs (VOI2) encompassing the entire calf muscle were drawn to determine background uptake. As the emission activity (A) inside VOI2 is composed of activity inside VOI1 and the background activity, A_VOI2_ = A_VOI1_ + A_Bg_, and activity is defined as the product of the uptake or tracer concentration (C) and the volume (V), A = C * V, the following formula can be derived:

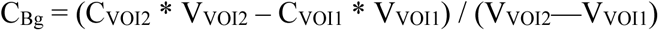

As activity inside VOI1 is the sum of activity inside the graft and background activity (A_VOI1_ = A_Graft_ + A_Bg_) and the graft volume (V_Graft_) was determined histologically, the uptake concentration inside the graft volume can be derived. These graft uptake values correct for dilution in uptake due to the enlarged VOI1 volume:

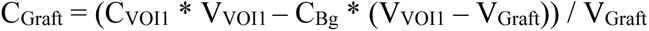

### Autoradiography

After the 3-month [^18^F]exendin or [^18^F]FDOPA PET scan, mice were sacrificed and their calf muscles excised and frozen in chilled isopentane and cut into 20-μm sections using a cryomicrotome (Microm HM 500 OM). The sections were exposed to an imaging plate (Fuji BAS Imaging Plate TR2025) for 4 hours and scanned using the Fuji BAS5000 analyzer. Images showing the tracer uptake were later compared to insulin immunohistochemistry of adjacent sections.

### Histological examination of graft volume and cyst proportion

After the final 5-month imaging timepoint, the mice were sacrificed, and their hind legs skinned. The calf muscles were harvested whole with a scalpel and fixed for 48h in 4% PFA in +25C followed by paraffin-embedding. These graft-bearing muscles were sectioned with a microtome until the graft was reached. From this first graft-containing section, 20-μm sections were taken while trimming until the whole graft was sectioned. Critically, depth trimmed and sectioned was recorded allowing calculation of total graft depth. Along the depth, at least 8 sections were stained with Hoechst separating nuclei-rich grafts and nuclei-sparse muscle; or analyzed unstained based on separation between autofluorescence-high muscle from autofluorescence-low grafts on the GFP channel. The section graft area was imaged on an EVOS microscope (Thermo Fisher) with 4x magnification and the images stitched using the “pairwise stitching” algorithm on Fiji (*60*). These stitched graft images were analyzed on a custom CellProfiler (version 4.0) (*61*) pipeline, in which the graft outlines were manually drawn and the area inside thresholded twice to include or exclude cysts, quantifying total and “cyst-free” graft areas in mm^2^. The sections divided the graft into parts. The volume of each part was calculated by multiplying the distance between the sections with the mean of section graft areas. Section area was defined as 0 for the first and last section containing graft tissue. The part volumes were summed for total graft volume.

### Histological examination of graft composition

Immunohistochemistry was conducted by running a standard deparaffinization series, followed by 30-minute HIER in +95C in sodium citrate buffer (pH 6), followed by 10-minute blocking (UltraV block, Thermo Fisher, TA-125-UB) and overnight application of primary antibodies in +4C. Secondary antibodies were applied for 45 minutes in +25C and slides mounted with ProLong antifade mountant (Thermo Fisher #P36984). The following primary antibodies were used: gp-insulin (Agilent/DAKO IR002, 1:2), mo-glucagon (Sigma-Aldrich #G2654, 1:500), mo-synaptophysin (Agilent/DAKO M7315, 1:200), rb-SLC18A1 (Sigma-Aldrich #HPA063797, 1:200), pan-specific sh-trypsin (R&D systems, #AF3586, 1:100), rb-cytokeratin-19 (Abcam 15463, 1:200). The following secondary antibodies were used, all with 1:500 dilution from Thermo Fisher: anti-mouse red (#21203), anti-mouse green (#21202), anti-rabbit green (#21206), anti-rabbit red (#21207), anti-guineapig red (#11076), anti-guineapig green (#11073) and anti-sheep green (#11015). Slides were imaged on Zeiss AxioImager equipped with Apotome II. Graft composition was analyzed from >10 random fields from 1-2 sections with a custom CellProfiler 4.0 pipeline, similar to one previously reported by our group (*53*). All images were analyzed so that all non-muscle areas were considered part of the graft

### Statistical methods

Prism version 9.5 (GraphPad software) was used for statistical analyses. Simple linear regression was conducted without forcing the lines to go through the origin, p-values given for slope being non-zero. Multiple linear regression model and correlation heatmaps were created with “Multiple Variables” feature on Prism. Welchs’s t-test was used when two groups were compared, and one or two-way ANOVA when there were more groups. No corrections were made for multiple comparisons, and data were assumed to be normally distributed. P-values were shorthanded as >0.05 ns, <0.05 *, <0.01 **, <0.001 ***.

## Acknowledgements

Vesa Oikonen (University of Turku) is thanked for advice regarding graft uptake calculations. Obada Al-Zghool (University of Turku) and Turku PET imaging center technicians Aake Honkaniemi, Marko Vehmanen, Mira Eisala and Päivi Kotitalo are thanked for assistance in PET image acquisition and processing, and processing of grafts. Solja Eurola (University of Helsinki) and Matthew Winder are thanked for genome editing. Onni Kolari (University of Eastern Finland) is thanked for assistance in graft sectioning, immunostaining and microscopy. Eliisa Vähäkangas, Vikash Chandra and Emilia Kuuluvainen (University of Helsinki) are thanked for feedback during project.

## Author contributions

Conceptualization: VL, TO, TG, KM, OS, PN

Genome editing: HI

SC-islet differentiation: VL

Implantation and animal follow-up: VL, HM, JSV, HI

Radiotracer synthesis: SL, LU, TK, OS, CBY, JR

Imaging: TG

Graft and SC-islet characterization: VL

Data analysis and interpretation: VL, TG, DB, TB

Funding: TO, PN, OS

Visualization: VL

Writing – original draft: VL, TO, TG

Writing – review and editing: VL, TB, TG, SL, DB, HM, TO, OS, PN

## Data and materials availability

All data are available upon reasonable request

