## Supplementary material for "Quantifying stem cell derived islet graft volume and composition with [^18^F]F-DBCO-exendin and [^18^F]FDOPA positron emission tomography": Figure supplements S1 to S6

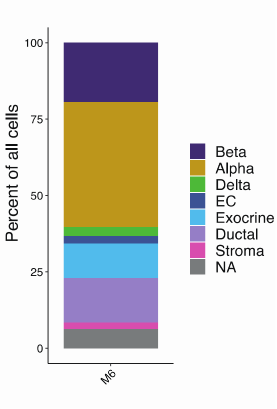


**Figure 1 supplement, cell type contributions to the single cell RNA sequencing:**
Kidey subcapsular SC-islet grafts at 6 months post-implantation, reanalyzed from *Balboa, Barsby & Lithovius 2022 Nature Biotechnology* data

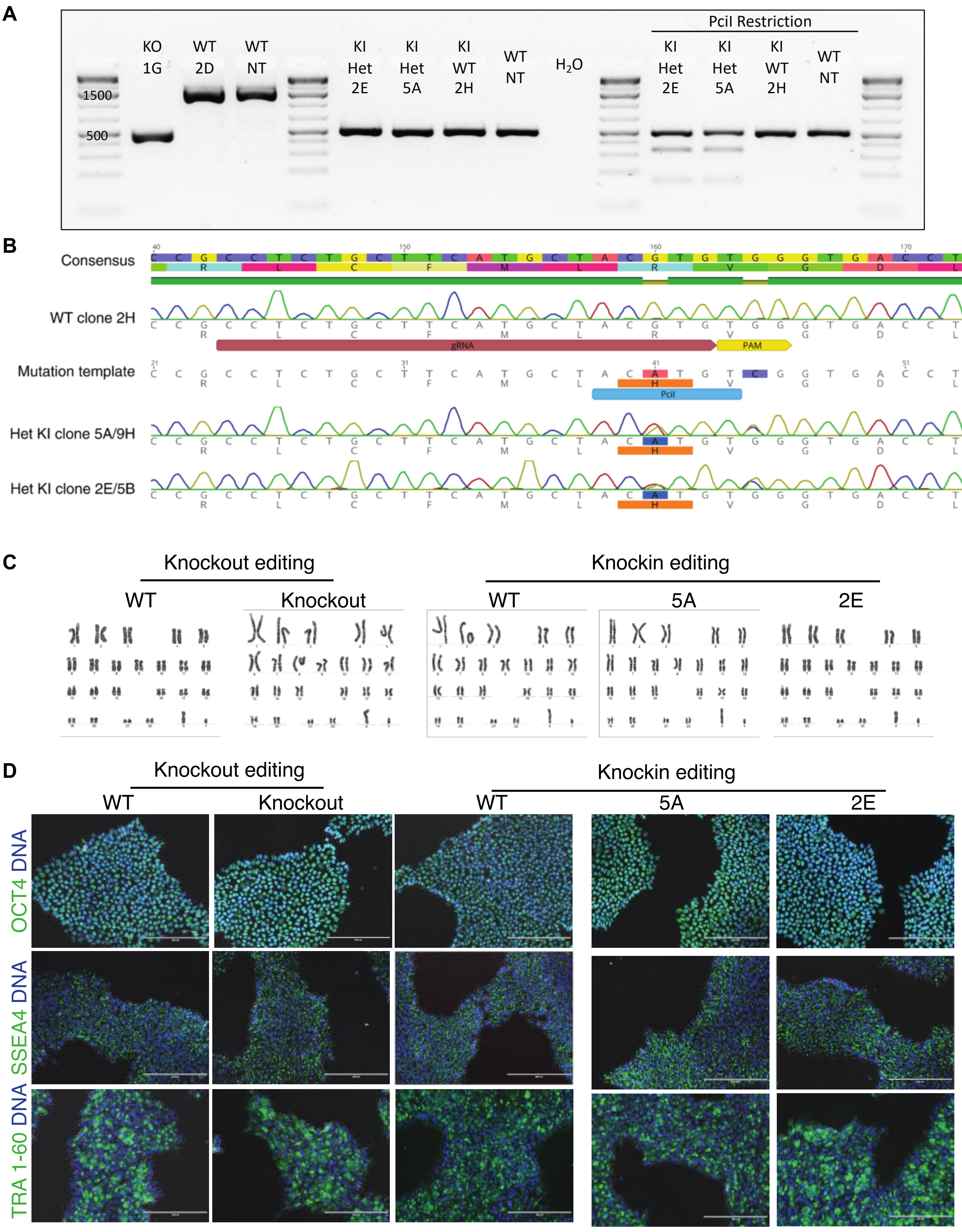


**Figure 2 supplement, quality control of KCNJ11 knockout and R201H knockin editing:**

**A**) PCR of the KCNJ11 locus of the knockouts (left) and of the knocked-in area containing PciI restriction site (right) **B**) Sanger sequencing of the knockin clones **C**) G-band karyotyping of the knockout and knockin clones **D**) Immunohistochemisty for pluripotency markers OCT4 (Santa Cruz, sc-9081, 1:250), SSEA4 (Thermo Fisher, MA1-021-D488, 1:100) and TRA 1-60 (Thermo Fisher, MA1-023, 1:100) in the knockout and knockin clones

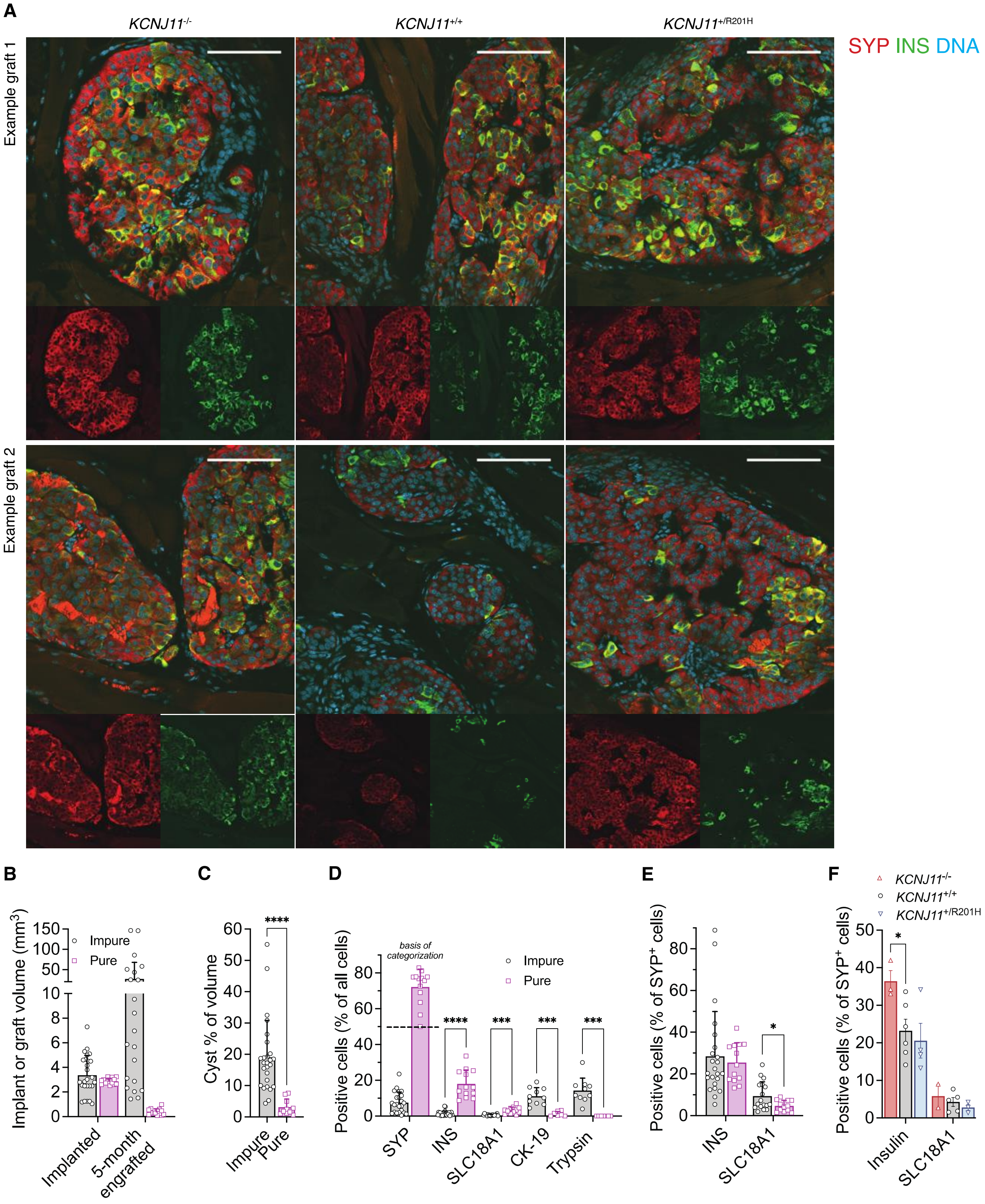


**Figure 3 supplement, composition data displayed by purity category and genotype:**

**A**) Example immunohistochemistry of pure 5-mo SC-islet grafts of different genotypes, stained for endocrine marker synaptophysin (SYP), beta cell marker insulin (INS) and nuclear marker hoechst, scale bar 100 $\mu m$ **B**) Implanted SC-islet volume and actual graft volume at 5-months post-implantation **C**) Cyst proportion of 5-month grafts **D**) Quantifications of immunohistochemisty, percentage of SYP+ served as basis for categorization to "pure" and "impure", cutoff at 50%. Enterochromaffin cell marker SLC18A1, exocrine marker trypsin and ductal marker cytokeratin-19 (CK-19) **E-F**) Quantification of INS+ and SLC18A1+ populations as percentage of SYP+ cells, grouped by purity E) and genotype F). All display mean±SD, one-way ANOVA or Welch's t-test.

u

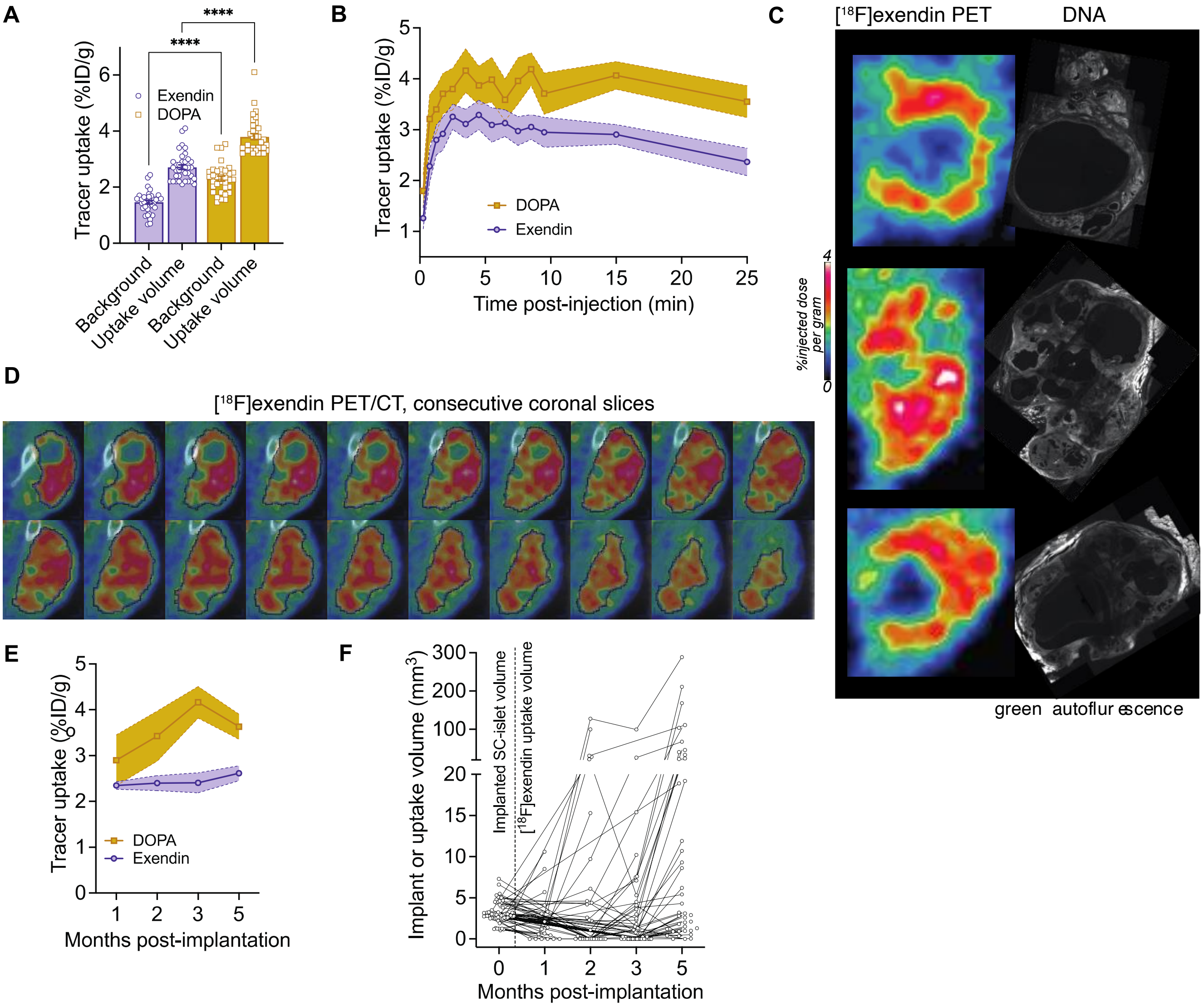


**Figure 4 supplement, additional PET data:**

**A**) Absolute levels of target and background tracer uptake at 5 months post-implantation (ratio in main figr e) , tw o - wa y ANOVA **B**) Dynamic uptake of the tracers during imaging at 5 months post-implantation as percentage of injected tracer dose per gram (=ml) of graft volume. Mean±95%CI, *N:* [18F]exendin=36, [18F]FDOPA=30, 25 min timepoint includes uptake from 20 to 30 minutes. **C-D**) Three distinct SC-islet grafts displaying low uptake areas inside the uptake volume corresponding to cysts in histology (top hoechst, middle and bottom green autoflur escence) C, and an addi tional graft where the uptake volume considered foreground (including cysts) is highlighted in 20 consecutive slices in PET/CT D), at 5 months post-implantation **E**) Progression of uptake concentration of both tracers during follow-up, mean±95%CI **F**) Graft size follow-up with [18F]exendin PET in individual grafts. Non-detected grafts marked with 0.

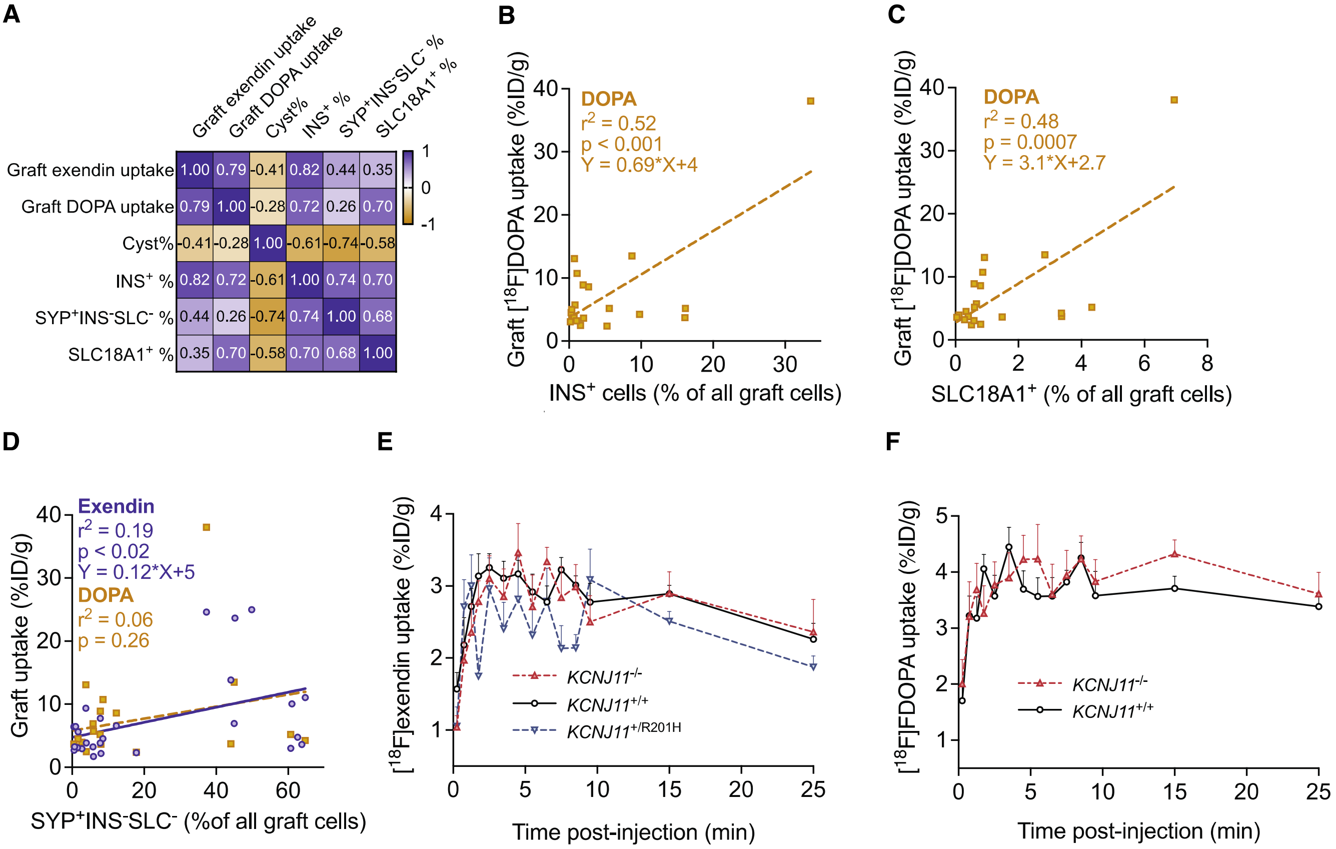


**Figure 5 supplement: Correlations of graft composition and genotype with tracer uptake:**

**A**) Pearson's correlation of the different composition parameters and graft tracer uptake **B-D)** Linear regression of graft uptake concentration and graft composition parameters: B) % of $INS^+$ beta cells C) % of $SLC18A1^+$ EC cell D) % of $SYP^+INS^-SLC18A1^-$ endocrine cells (i.e. alpha and delta cells) of all graft cells **E-F**) Dynamic uptake of [$^{18}$F]exendin E) and [$^{18}$F]DOPA F) during imaging at 5 months post-implantation in cohort-2, each genotype separate. A single *KCNJ11*$^{+/R201H}$ graft was detected with DOPA and is not plotted

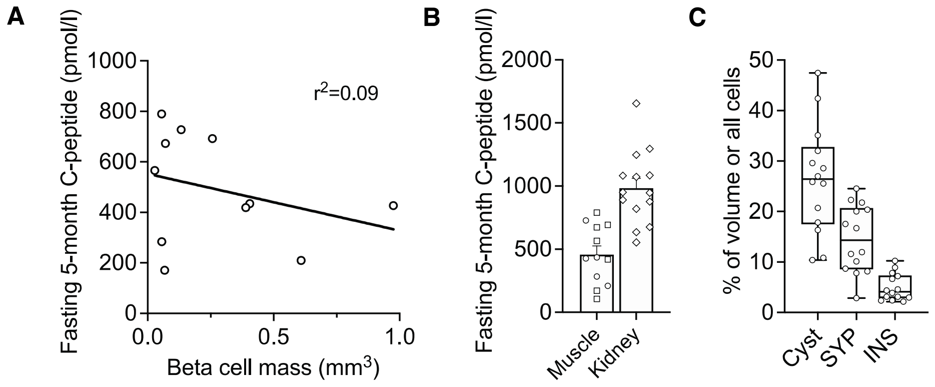


**Figure 6 supplement, intramuscular graft C-peptide measurements and kidney graft composition:**
**A**) Linear regression of the circulating C-peptide at 5 months and the sum of the non-cystic, beta cell fraction corrected graft volumes in the two intramuscular grafts in the mice used for the imaging studies. **B**) Fasting C-peptide in the mice used for imaging (muscle, the mice carry two grafts, cohorts pooled) and in thekidney cohort at 5-months post-implantation **C**) Cystic volume as percentage of total graft volume, and the percentage of $SYP^+$ endocrine cell and $INS^+$ beta cells out of all cells in the graft (including connective tissue cells)
